# A causal role for right frontopolar cortex in directed, but not random, exploration

**DOI:** 10.1101/064741

**Authors:** Wojciech Zajkowski, Malgorzata Kossut, Robert C. Wilson

## Abstract

The explore-exploit dilemma occurs anytime we must choose between exploring unknown options for information and exploiting known resources for reward. Previous work suggests that people use two different strategies to solve the explore-exploit dilemma: directed exploration driven by information seeking and random exploration driven by decision noise. Here, we show that these two strategies rely on different neural systems. Using transcranial magnetic stimulation to selectively inhibit right frontopolar cortex, we were able to selectively inhibit directed exploration while leaving random exploration intact, suggesting a causal role for right frontopolar cortex in directed, but not random, exploration.

## Results and Discussion

In an uncertain world, adaptive behavior requires us to carefully balance the exploration of new opportunities with the exploitation of known resources. Finding the optimal balance between exploration and exploitation is a hard computational problem and there is considerable interest in how humans and animals strike this balance in practice (Badre et al, 2012; Cavanagh et al, 2011; Cohen et al, 2007; Daw et al, 2006; Frank et al, 2009; Hills et al, 2015; Mehlhorn et al, 2015, Wilson et al, 2014). Recent work has suggested that humans use two distinct strategies to solve the explore-exploit dilemma: directed exploration, based on information seeking, and random exploration, based on decision noise (Wilson et al, 2014). Even though both of these strategies serve the same purpose, i.e. balancing exploration and exploitation, it is likely they rely on different cognitive mechanisms. Directed exploration is driven by information and is thought to be computationally complex (Wilson et al, 2014). On the other hand, random exploration can be implemented in a simpler fashion by using neural or environmental noise to randomize choice.

A reasonable, biologically-guided assumption is that these two types of exploration rely on separate brain mechanisms. Of particular interest is the right frontopolar cortex (RFPC) an area that has been associated with a number of functions, such as tracking alternate options (Boorman et al, 2009), strategies (Domenech & Koechlin, 2015) and goals (Pollmann, 2015) that may be important for exploration. In addition, a number of studies have implicated the frontal pole in exploration itself, although importantly, how exploration is defined varies from paper to paper. In one line of work, exploration is defined as information seeking. Defined this way, exploration correlates with FP activity measured via fMRI (Badre et al, 2012) and a frontal theta component in EEG (Cavanagh et al, 2011), suggesting a role for FP in *directed* exploration. However, in another line of work, exploration is defined differently, as choosing the low value option, not the most informative. Such a measure of exploration is more consistent with *random* exploration where decision noise drives the sampling of low value options by chance. Defined in this way, exploratory choice correlates with FP activation (Daw et al, 2006) and stimulation and inhibition of FP with direct current (tDCS) can increase and decrease the frequency with which such exploratory choices occur (Beharelle et al, 2015).

Taken together, these two sets of findings suggest that FP plays a crucial role in both directed *and* random exploration. However, we believe that such a conclusion is premature because of a subtle confound that arises between reward and information in most explore-exploit tasks. This confound arises because participants only gain information from the options they choose, yet they are incentivized to choose more rewarding options. Thus, over many trials, participants gain more information about more rewarding options such that the two ways of defining exploration, choosing high information or low reward options, become confounded (Wilson et al, 2014). This makes it impossible to tell whether the link between FP and exploration is specific to either directed or random exploration, or whether it is general to both.

To distinguish these interpretations and investigate the causal role of RFPC in directed and random exploration, we used continuous theta-burst TMS (Huang et al, 2005) to selectively inhibit RFPC in fifteen participants performing the “Horizon Task”, an explore-exploit task specifically designed to separate directed and random exploration (Wilson et al, 2014). Using this task we find evidence that RFPC inhibition selectively inhibits directed exploration while leaving random exploration intact.

We used our previously published “Horizon Task” (Figure 1) to measure the effects of TMS stimulation to RFPC on directed and random exploration. In this task, participants play a set of games in which they make choices between two slot machines (one-armed bandits) that pay out rewards from different Gaussian distributions. To maximize their rewards in each game, participants need to exploit the slot machine with the highest mean, but they cannot identify this best option without exploring both options first.

**Figure 1.**
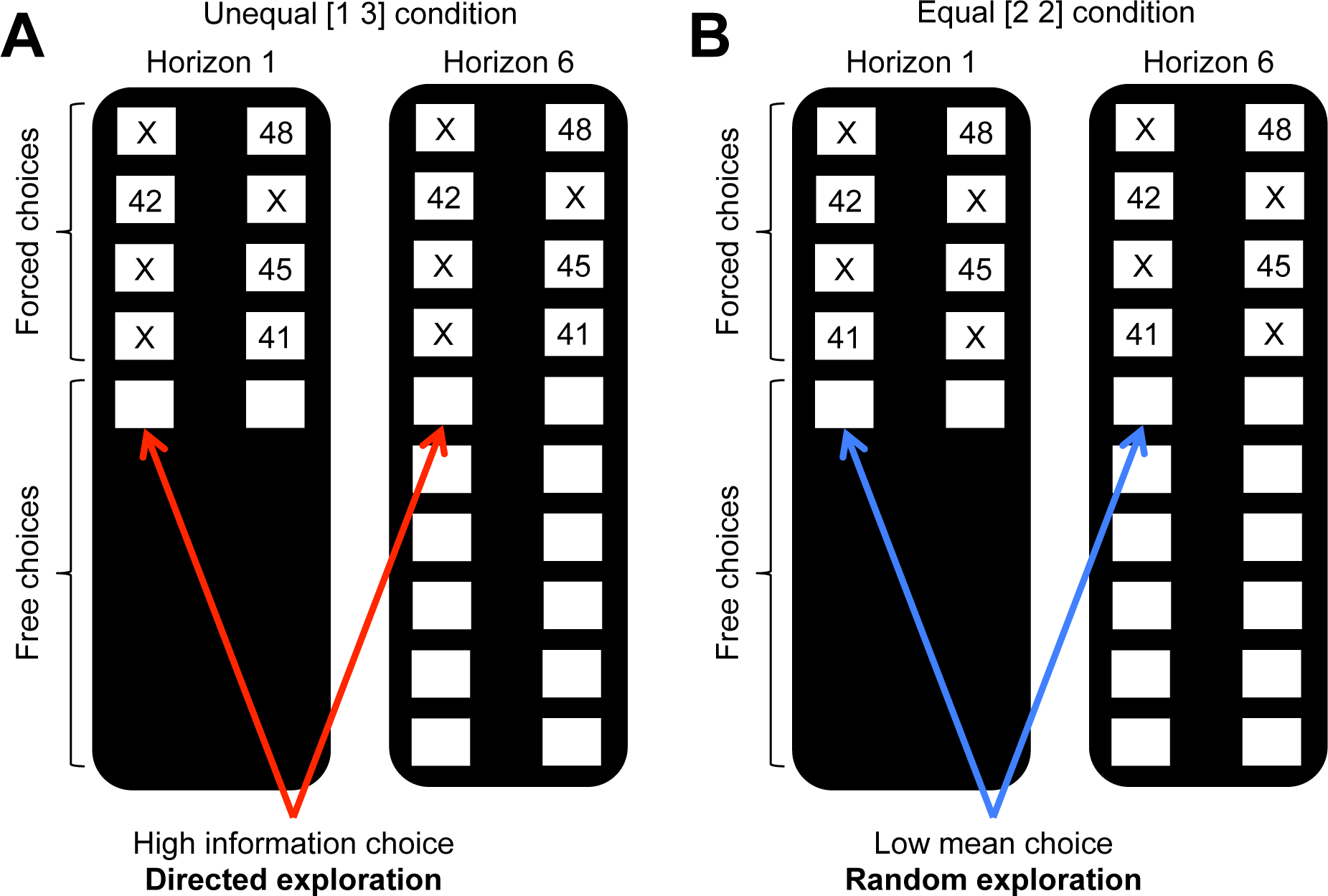
The Horizon Task. Participants make a series of decisions between two one-armed bandits that pay out probabilistic rewards with unknown means. At the start of each game, forced-choice trials give participants partial information about the mean of each option setting up one of two information conditions: (**A**) an unequal (or [1 3]) condition in which participants see 1 play from one option and 3 plays from the other and (**B**) an equal (or [2 2]) condition in which participants see 2 plays from both options. Directed exploration is then defined as the change in information seeking with horizon in the unequal condition (**A**). Random exploration is defined as the change choosing the low mean option in the equal condition (**B**).

The key manipulation in the Horizon Task is the time horizon, the number of decisions remaining in each game. The idea behind this manipulation is that when the horizon is long (6 trials), participants should explore, because any information they acquire can be used to make better choices later on. In contrast, when the horizon is short (1 trial), participants should exploit the option they believe to be best. Thus, by measuring *changes* in information seeking and behavioral variability that occur with horizon, this task allows us to quantify directed and random exploration.

The Horizon Task also allows us to remove the reward-information confound with the use of “forced-choice” trials at the start of each game. These forced-choice trials setup one of two information conditions: an unequal (or [1 3]) condition (Figure 1A), in which one option is played once and the other three times, and an equal (or [2 2]) condition (Figure 1B), in which both options are played twice. Information seeking is quantified as the probability of choosing the more uncertain option in the [1 3] condition (i.e. the option played once in the forced-choice trials), p(high info). Behavioral variability is quantified as the number of mistakes, i.e. choosing the low value option, in the [2 2] condition, p(low mean). To remove the reward-information confound, both of these measures are computed on the *first* free-choice trial in each game, i.e. before participants’ choices have a chance to confound information and reward.

Using these measures of exploration, we found that inhibiting the RFPC had a significant effect on directed exploration but not random exploration (Figure 2A, B). In particular, for directed exploration, a repeated measures ANOVA revealed a significant interaction between stimulation condition and horizon (F(1, 14) = 4.77, p = 0.047). Conversely, a similar analysis for random exploration revealed no effects of stimulation condition (main effect of stimulation condition, F(1, 14) = 0.26, p = 0.62; interaction of stimulation condition with horizon, F(1, 14) = 0.01; p = 0.93). Post hoc analyses revealed that the change in directed exploration was driven by changes in information seeking in horizon 6 (one-sided t-test, t(14) = 2.26, p = 0.02) and not in horizon 1 (two-sided t-test, t(14) = -0.40, p = 0.69). Finally, a similar analysis using a logistic model of choice behavior yielded similar findings (see Supplementary Material).

**Figure 2.**
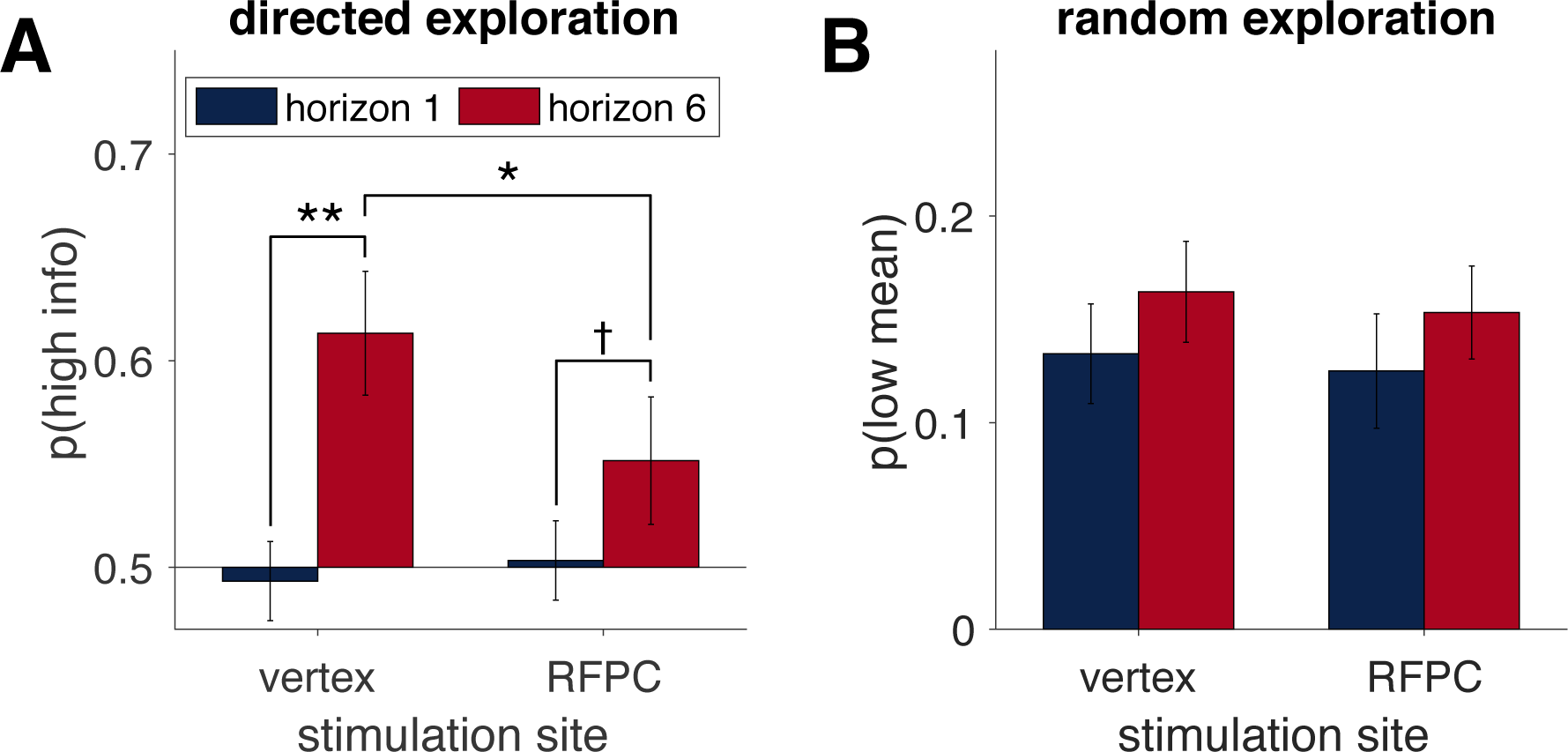
RPFC stimulation affects directed, but not random, exploration. (**A**) In the control (vertex) condition, information seeking increases with horizon, consistent with directed exploration. When RFPC is stimulated, directed exploration is reduced, an effect that is entirely driven by changes in horizon 6 († denotes p < 0.1, * p < 0.05, ** p < 0.01). (**B**) Random exploration increases with horizon but is not affected by RFPC stimulation.

These results suggest a causal role for RFPC in directed, but not random, exploration. A role in directed exploration is consistent with other findings implicating FP in a number of complex computations. These include tracking the value of the best unchosen option (Boorman et al, 2009), inferring the reliability of alternate strategies (Domenech & Koechlin, 2015), arbitrating between old and new strategies (Donoso et al, 2015; Mansouri et al, 2015), and reallocating cognitive resources among potential goals in underspecified situations (Pollmann, 2015). Taken together, these findings suggest a role for frontal pole in decisions that involve long-term planning and the consideration of alternative actions, both crucial for directed exploration.

That frontal pole is *not* involved in random exploration suggests that directed and random exploration rely on (at least partially) dissociable neural systems. Exactly what these systems are is currently unknown, but may include other areas of prefrontal cortex (Badre et al, 2012; Cavanagh et al, 2011; Daw et al, 2006) and may involve modulation by noradrenergic (Cohen et al, 2007) and dopaminergic (Costa et al, 2014) inputs. Clearly more work, involving tasks that can dissociate the two types of exploration, will be required to fully understand the neural substrates of exploratory choice.

## Methods

### Participants

16 healthy right-handed, adult volunteers (9 female; aged 19-32). One participant (female) was excluded from the analysis due to chance-level performance in both experimental sessions. All participants were informed about potential risks connected to TMS and signed a written consent. The study was approved by University of Social Sciences and Humanities ethics committee.

### Procedure

There were two experimental TMS sessions and a preceding MRI session. On the first session T1 structural images were acquired using a 3T Siemens TRIO scanner. The scanning session lasted up to 10 minutes. Before the first two sessions, participants filled in standard safety questionnaires regarding MRI scanning and TMS. During the experimental sessions, prior to the stimulation participants went through 16 training games to get accustomed to the task. Afterwards, resting motor thresholds were obtained and the stimulation took place. Participants began the main task immediately after stimulation. The two experimental sessions were performed with an intersession interval of at least 5 days. The order of stimulation conditions was counterbalanced across subjects. All sessions took place at Nencki Institute of Experimental Biology in Warsaw.

### Stimulation site

The RFPC peak was defined as [x,y,z]= [35,50,15] in MNI (Montreal Neurological Institute) space. The coordinates were based on a number of fMRI findings that indicated RFPC involvement in exploration^1,7^ and constrained by the plausibility of stimulation (e.g. defining ‘z’ coordinate lower would result in the coil being placed uncomfortably close to the eyes). Vertex corresponded to the Cz position of the 10-20 EEG system. In order to locate the stimulation sites we used a frameless neuronavigation system (Brainsight software, Rogue Research, Montreal, Canada) with a Polaris Vicra infrared camera (Northern Digital, Waterloo, Ontario, Canada).

### TMS protocol

We used continuous theta burst stimulation (cTBS)^13^. cTBS requires 50Hz stimulation at 80% resting motor threshold. 40 second stimulation is equivalent to 600 pulses and can decrease cortical excitability for up to 50 minutes (Wischnewski & Schutter, 2015). Individual resting motor thresholds were assessed by stimulating the right motor knob and inspecting if the stimulation caused an involuntary hand twitch in 50% of the cases. We used a MagPro X100 stimulator (MagVenture, Hueckelhoven, Germany) with a 70mm figure-eight coil. The TMS was delivered in line with established safety guidelines (Rossi et al, 2009).

### Limitations

Defining stimulation target by peak coordinates based on findings from previous studies did not allow to account for individual differences in either brain anatomy or the impact of TMS on brain networks (Gratton et al, 2013). However, a study by Volman and collegues (2011) that used the same theta-burst protocol on the left frontopolar cortex has shown biletaral inhibitory effects on blood perfusion in the frontal pole. This suggests that both right and left parts of the frontopolar cortex might have been inhibited in our experiment, which is consistent with imaging results indicating bilateral involvement of the frontal pole in exploratory decisions.

### Task

The task was a modified version of the Horizon Task (Wilson et al, 2014). As in the original paper, the distributions of payoffs tied to bandits were independent between games and drawn from a Gaussian distribution with variable means and fixed SD=8. Participants were informed that in every game one of the bandits is objectively ‘better’ (has a higher payoff mean). Differences between two means were set to either 4, 8, 12 or 20. One of the means was always equal to either 40 or 60 and the second was set accordingly. The order of games was randomized. Mean sizes and order of presentation were counterbalanced. Participants played 160 games and the whole task lasted between 39 and 50 minutes (m=43.4).

Each game consisted of 5 or 10 choices. Every game started with a screen saying “New game” and information about whether it was a long or short horizon, followed by sequentially presented choices. Every choice was presented on a separate screen, so that participants had to keep previous the scores in memory. There was no time limit for decisions. During forced choices participants had to press the prompted key to move to the next choice. During free choices they could press either ‘z’ or ‘m’ to indicate their choice of left or right bandit. The decision could not be made in a time shorter than 200ms, preventing participants from accidentally responding too soon. The score feedback was presented for 500ms. A counter at the bottom of the screen indicated the number of choices left in a given game. The task was programmed using PsychoPy software v1.86 (Peirce, 2007).

Participants were rewarded based on points scored in two sessions. The payoff bounds were set between 50 and 80 zl (equivalent to approximately 12 and 19 euro). Participants were informed about their score and monetary reward after the second session.

## Supplementary Material

### Model-based analysis

To complement our analysis in the main paper, we used a model-based analysis based on the model described in Wilson et al. (2014). Briefly, this model assumes that participants make their decision on the first free-choice trial based on the difference in the observed mean reward between left and right bandits, *Δμ*, and the difference in information between left and right bandits, *ΔI* (= +1 when left is more informative in the [1 3] condition, -1 when the right option is more informative in the [1 3] condition and 0 in the [2 2] condition). In particular, we assume that participants choose the left option with probability *p_left_* which is given by

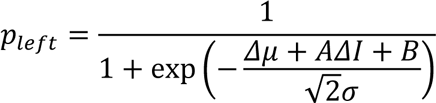

where *A* is the information bonus that quantifies the value of information and directed exploration, *σ* is the standard deviation of the decision noise that quantifies random exploration, and *B* is the spatial bias that accounts for any baseline tendency to favor left or right choices.

As described in Wilson et al. (2014), these three parameters, *A*, *σ* and *B*, were fit separately for each subject in each horizon and information condition using a maximum *a posteriori* approach. In line with our previous work, we assumed a Gaussian prior with mean zero and standard deviation 20 on *A*, an exponential prior with length scale 20 for *σ* and no prior on *B*.

In line with our model-free analysis in the main text, we found that TMS stimulation of RFPC had a significant effect on directed, but not random, exploration (Figure S1). For directed exploration, a repeated measures ANOVA revealed a significant interaction between horizon and stimulation condition (F(1,14) = 5.11 p = 0.04). For random exploration in the [2 2] condition there was no such interaction (F(1, 14) = 0.93, p = 0.35). In addition to measuring decision noise in the [2 2] condition, the model-based analysis also allows us to measure random exploration in the [1 3] condition. Again we found no interaction of horizon and condition (F(1, 14) = 0.93, p = 0.35) further bolstering our claim that RFPC has no effect on random exploration. Post hoc analysis of the directed exploration result was also consistent with the model-free findings in that we found that the interaction effect was entirely driven by changes in horizon 6 (one-sided t-test t(14) = 2.21, p = 0.022) not horizon 1 (two-side t-test, t(14) = -0.50, p = 0.63).

**Figure S1.**
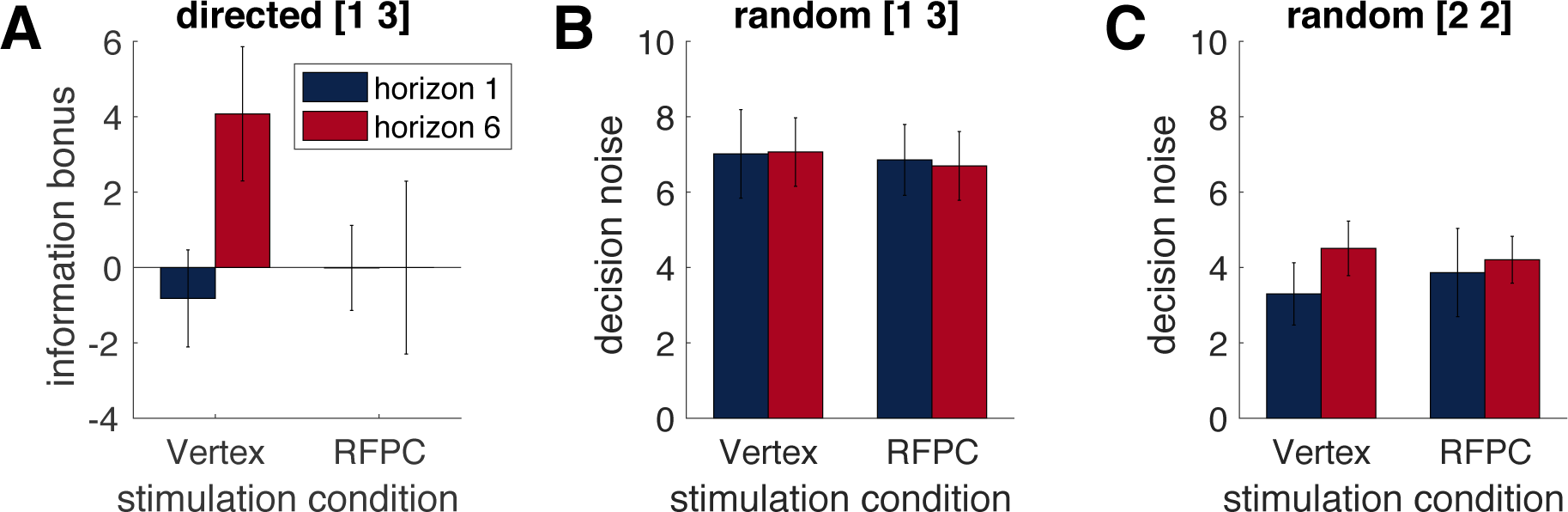
Mean parameter values for the model-based analysis.

Finally, as another way to visualize the result and to gain a qualitative sense of the quality of the model fit, we computed choice curves for each stimulation condition in the [1 3] condition (Figure S2). These choice curves plot p(high info) as a function of the difference in mean between the high and low information options. In these plots, directed exploration manifests as a left-shift of the horizon 6 curve relative to the horizon 1 curve, as clearly seen in the control condition (Figure S2A). When RFPC is stimulated, this left-shift of the choice curve disappears (Figure S2B) consistent with our other results. These plots also show relatively good agreement between the empirical choice curves computed directly from the data and the fit choice curves computed from the model.

**Figure S2.**
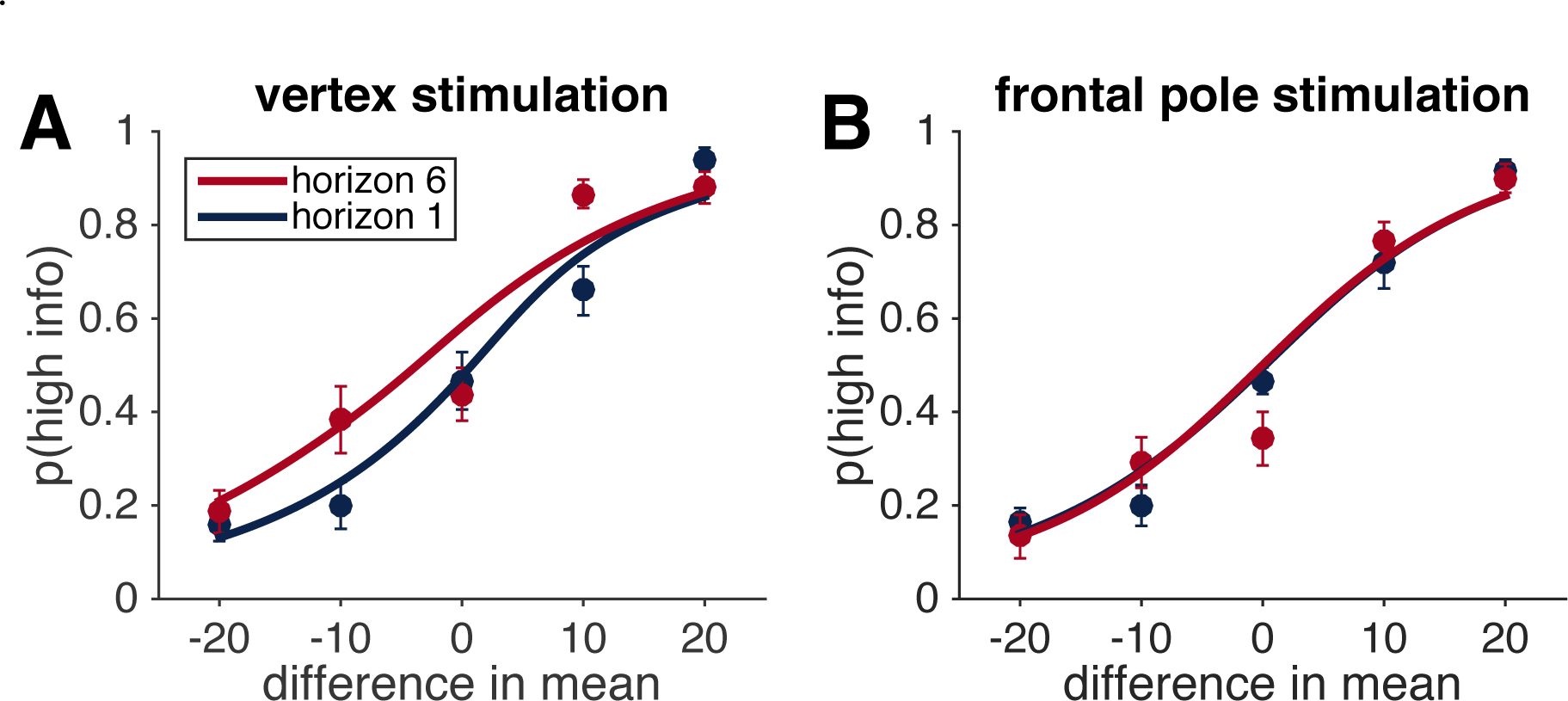
**Choice curves** showing p(high info) as a function of the difference in mean between the more and less informative options in the [1 3] condition.

